# TrkB Agonist LM22A-4 Increases Oligodendroglial Populations During Myelin Repair in the Brain

**DOI:** 10.1101/642918

**Authors:** Huynh TH Nguyen, Rhiannon J Wood, Alexa R Prawdiuk, Sebastian GB Furness, Junhua Xiao, Simon S Murray, Jessica L Fletcher

## Abstract

The neurotrophin, brain-derived neurotrophic factor (BDNF) promotes central nervous system (CNS) myelination during development and after injury. This is achieved via activation of oligodendrocyte-expressed tropomyosin-related kinase (Trk) B receptors. However, while administration of BDNF has shown beneficial effects, BDNF itself has a poor pharmacokinetic profile. Here, we compare two TrkB-targeted BDNF-mimetics, the structural-mimetic, tricyclic dimeric peptide-6 (TDP6) and the non-peptide small molecule TrkB agonist LM22A-4 in the cuprizone model of central demyelination in female mice. Both mimetics promoted remyelination, increasing myelin sheath thickness and oligodendrocyte densities after one-week recovery. Importantly, LM22A-4 exerts these effects in an oligodendroglial TrkB-dependent manner. However, analysis of TrkB signaling by LM22A-4 suggests rather than direct activation of TrkB, LM22A-4 exerts its effects via indirect transactivation of Trk receptors. Overall, these studies support the therapeutic strategy to selectively targeting TrkB activation to promote remyelination in the brain.

## 1 Introduction

The neurotrophin, brain-derived neurotrophic factor (BDNF) is an attractive therapeutic for many neurodegenerative diseases due to its broad neuroprotective effects promoting neuronal survival, synaptic plasticity and central nervous system (CNS) myelination (Chao, 2003; Fletcher et al., 2018b; Longo and Massa, 2013). Its action *via* oligodendrocyte expressed TrkB to potentiate and enhance myelination (Du et al., 2003; Wong et al., 2013; Xiao et al., 2011) makes it particularly promising for central demyelinating diseases such as multiple sclerosis (MS), where there is an unmet clinical need for remyelinating therapies to halt disease progression. However, BDNF itself has poor pharmacokinetic properties; it is non-selective, also acting through the pan-neurotrophic receptor p75^NTR^, has a short-half life and has high molecular weight, limiting its ability to penetrate the blood-brain barrier (Longo and Massa, 2013; Poduslo and Curran, 1996). To overcome these limitations a range of small molecule BDNF-mimetics that selectively target the TrkB receptor have been developed (Longo and Massa, 2013). This includes tricyclic dimeric peptide-6 (TDP6) (O’Leary and Hughes, 2003) and the partial TrkB agonist, LM22A-4 (Massa et al., 2010).

TDP6 is a structural peptide mimetic, designed to mimic the Loop 2 region of BDNF that is known to interact with TrkB (Chao, 2003; O’Leary and Hughes, 2003). We have previously shown that TDP6 mimics BDNF in promoting neuronal survival (O’Leary and Hughes, 2003) and enhancing myelination both *in vitro* (Wong et al., 2014) and during myelin repair following cuprizone demyelination *in vivo* (Fletcher et al., 2018a). Similarly, LM22A-4 was identified during an *in silico* screen to identify compounds with potential to mimic the Loop2 region of BDNF and is a non-peptide, partial TrkB agonist (Massa et al., 2010). It is 98% smaller than BDNF and has been shown to have therapeutic potential in preventing neurodegeneration in animal models of traumatic brain injury, stroke, Huntington’s disease and Rhett syndrome (Gu et al., 2018; Han et al., 2012; Massa et al., 2010; Schmid et al., 2012; Simmons et al., 2013) but has not yet been tested in the context of central demyelinating disease such as MS.

Here, we compare the effect of intra-cerebroventricular (ICV) administration of these BDNF mimetics following cuprizone demyelination in mice. Both mimetics promoted remyelination, in particular myelin sheath thickness, after one-week recovery. Interestingly, LM22A-4 increased the density of oligodendroglia in the corpus callosum more than TDP6. Importantly, these effects were dependent on TrkB, as post-cuprizone treatment with LM22A-4 in mice with conditional deletion of TrkB from oligodendrocytes abrogated the effects on both remyelination and oligodendroglial density. While this indicates that LM22A-4 promotes myelin repair in a TrkB dependent manner, assessment of TrkB phosphorylation and signaling *in vitro* suggests that LM22A-4 may not activate TrkB directly, but rather result in delayed TrkB transactivation *via* a GPCR-mediated mechanism. Collectively these data further verify that targeting TrkB activation is a cogent strategy to promote myelin repair in the brain, and that alternate small molecule mimetic strategies are effective towards this end. Further studies aimed at elucidating the precise mechanism of action are warranted to optimize the therapeutic potential of this approach.

## 2 Materials and Methods

### 2.1 Experimental animals and cuprizone induced demyelination

Female C57BL/6 mice aged 8 weeks were fed 0.2% cuprizone in normal chow (Teklad Custom Research Diets, USA) for 6 weeks to induce demyelination. Cuprizone feed was removed, and mice were sacrificed or received intracerebroventricular osmotic pumps for 7 days.

For experiments in conditional knockout mice, female, 8-10 week-old CNPase^+/−^ x TrkB^fl/fl^ (Fletcher et al., 2018a; Lappe-Siefke et al., 2003; Lulkart et al., 2005) on C57BL/6 background underwent the procedures described above.

All mice were housed in specific pathogen free conditions at the Melbourne Brain Centre Animal Facility. All animal procedures were approved by the Florey Institute for Neuroscience and Mental Health Animal Ethics Committee and followed the Australian Code of Practice for the Care and Use of Animals for Scientific Purposes.

### 2.2 Intracerebroventricular delivery of LM22A-4 and TDP6

Following cuprizone feeding, mice received either: 40µM TDP6, 500µM LM22A-4 or the artificial cerebrospinal fluid (aCSF) vehicle intracerebroventricularly (ICV) as described in Fletcher et al., 2018a. Briefly, cannulae of osmotic pumps (flow rate: 0.5µL/hr, Azlet) were stereotaxically inserted over the right lateral ventricle under isoflurane anesthesia. Infusion concentrations of TDP6 and LM22A-4 were determined based on previous *in vitro* studies characterizing their effective concentrations (Massa et al., 2010; Wong et al., 2014). TDP6 infused animals were the same animals from Fletcher et al. 2018 and LM22A-4 infusion experiments were performed concurrently. Following stereotaxic surgery all mice were placed in a recovery chamber maintained at 32°C and were monitored for adverse reactions immediately following surgery and then daily. After 7 days of ICV infusion, mice were taken for necropsy and brain removed for immunostaining and electron microscopy (EM).

### 2.3 Tissue processing and immunofluorescence

Mice were anaesthetized and transcardially perfused with 0.1M sterile mouse isotonic phosphate buffered saline (PBS) followed by 4% paraformaldehyde (PFA). Brains were collected and post-fixed overnight in 4% PFA. The first millimeter of the right hemisphere from the sagittal midline was selected for EM processing as previously described (Fletcher et al., 2018a). The remaining tissue and contralateral hemisphere were cryoprotected in 30% sucrose prior to embedding in OCT. Frozen sections were cut in the sagittal orientation at 10µm thickness using a cryostat maintained between −20 to −17°C and collected on SuperfrostPlus slides, air-dried and stored at −80°C until use. Approximately 70-100µm separated adjacent sections on each slide.

Immunofluorescent staining was performed as previously described (Fletcher et al., 2018a). Briefly, slides were washed in PBS before overnight incubation at room temperature with primary antibodies. Slides were then washed and incubated with the appropriate fluorophore-conjugated secondary antibody for 2 hours at room temperature in the dark. Slides were washed, and counterstained with nuclear marker Hoescht33442 before mounting with aqueous mounting media (DAKO). All immunohistochemistry was performed in batches.

Antibodies used were: rat anti-myelin basic protein (MBP, 1:200, MAB386, Millipore, MA, USA), rabbit anti-Olig2 (1:200, AB9610, Millipore, MA, USA), mouse anti-CC1 (1:200, APC, OP80, CalBioChem, CA, USA), goat anti-platelet derived growth factor receptor-α (PDGFRα, 1:200, AF1062. R&D Systems, MN, USA), goat anti-Iba1 (1:200, ab5076, Abcam, UK), mouse anti-glial fibrially acidic protein (GFAP, 1:100, MA360, Millipore, MA, USA) and rabbit anti-phosphorylated TrkB^S478^ (1:200 R-1718-50, Biosensis).

### 2.4 Electron microscopy and analysis

Semi-thin (0.5-0.1µm) sections of caudal corpus callosum in a sagittal plane were collected on glass slides and stained with 1% toluidine blue to select region of analysis. Ultrathin (0.1µm) sections were subsequently collected on 3×3mm copper grids and specimens examined using a JEOL 1001 transmission electron microscope. Images were captured with MegaView III CCD cooled camera operated with iTEM AnalySIS software (Olympus Soft Imaging Systems GmbH). A minimum of six distinct fields of view were imaged at 5000 or 10000x magnification for each animal. The proportion of myelinated axons, axon diameter and g-ratio were analysed manually using FIJI/ImageJ (National Institutes of Health). For g-ratios at least 100 axons from 3 mice per group were measured. Resin embedding, sectioning and post-staining and EM imaging were performed at the Peter MacCallum Centre for Advanced Histology and Microscopy.

### 2.5 Fluorescence imaging and analysis

Imaging was performed blind to treatment group and restricted to the caudal corpus callosum approximately −1.1 to −3.0mm from Bregma. Tracts contributing to the dorsal hippocampal commissure were excluded from analysis. For each analysis, a minimum of three sections per animal were imaged.

To quantify the level of remyelination images of MBP stained sections were collected with an AxioVision Hr camera attached to a Zeiss Axioplan2 epi-fluoresence microscope under a 20x objective. Uniform exposure times were used. Remaining images were acquired with a Zeiss LSM780 or LSM880 confocal microscope with 405nm, 488nm, 561nm and 633nm laser lines. For each fluorescent stain uniform settings were used.

MBP staining was measured as described in Fletcher et al. 2014 using the threshold function in FIJI/Image J and limited to a standard region of interest (ROI) of 625000µm^2^ for each section. Data were expressed as a percentage area of positive staining in a single ROI.

#### 2.5.1 Cell counts

All cell counts were performed blind to sample identity, manually in FIJI/Image J. Data were expressed as the number of cell/mm^2^ or proportion out of the total number of nuclei.

### 2.6 Generation of isogenic TrkB expressing Flp-In 293 cells

The Flp-In 293 cell system (ThermoScientific) was used to generate isogenic TrkB expressing HEK293 cell lines where TrkB was integrated into the host cell genome. Flp-In cells express a hygromycin resistance gene, which enables the use of hygromycin (50µg/µL) to exert selective pressure for cells carrying the Flp-In construct.

Briefly, to generate the TrkB expressing HEK293 cell line (Figure S1A), Ntrk2 (NM_012731.2) was amplified by PCR from rat cDNA and recombined into pDONR201 entry vector by BP clonase II reaction (Invitrogen) according to the manufacturer’s instructions. DH5α bacteria (ThermoScientific) were transformed with the entry vector by heat shock and positive colonies expressing the pDONR201 plasmid were selected using kanamycin resistance. pDONR201 plasmid containing Ntrk2 was purified using a mini-prep kit (Promega) and recombined into pEF5/FRT/V5-DEST destination vector by LR clonase II reaction (ThermoScientific). Following transformation with the destination vector, DH5α bacterial colonies were placed under ampicillin selective pressure and plasmid DNA extracted. At each step, successful recombination of Ntrk2 into the entry and destination vectors was confirmed by restriction ligase digest with ApaI (NEB) and Sanger sequencing (Australian Genome Research Facility).

Once the Ntrk2 destination vector was generated, the Flp-In HEK293 host cells were transfected with the Ntrk2 destination vector and the Flp-recombinase vector pOG44 with 50µg/µL hygromycin according to the manufacturer’s instructions. Flp-In HEK293 cells were maintained at 37°C with 5% CO2 in Dulbecco’s modified eagle medium (DMEM) with 10% fetal bovine serum, 1% L-glutamine, 1% penicillin and 1% streptomyocin. TrkB expression by the isogenic TrkB Flp-In HEK293 cells was verified by Western blot (Figure S1B-E).

### 2.7 *In vitro* testing of TrkB phosphorylation by LM22A-4

To examine the capacity of LM22A-4 to mimic the BDNF-TrkB signaling cascade, isogenic TrkB FlP-In HEK293 cells were starved in serum free media for 2 hours before treatment with 4nM BDNF or 500nM LM22A-4 for 0, 5, 15, 30, 60 and 240mins. Concentrations were chosen based on previous work (Massa et al., 2010; Wong et al., 2014). Cells were lysed with TNE (Tris) buffer containing protease (Complete Mini) and phosphatase inhibitors (PhosStop Roche, 50mM Sodium Fluoride). Protein concentrations were determined by Bradford assay and lysates stored at −80°C until use.

### 2.8 SDS-PAGE and Western blot analysis

Lysates were separated by SDS-PAGE (4-12% Bis-Tris, Invitrogen) and transferred to PVDF membrane and probed with antibodies against TrkB (1:1000, sc-8316, SantaCruz) and pTrkB^s478^ (1:1000, R-1718-50, Biosensis), p44/42 MAPK (ERK1/2, 1:1000, #9102 Cell Signaling Technologies) and phosphorylated ERK1/2 (pERK1/2, 1:1000, #9101 Cell Signaling Technologies). All blots shown are representative of at least 3 independent experiments. Optical density value for each band was determined using FIJI/ImageJ and corrected to loading control and normalized against the relevant control condition.

### 2.9 Statistical analyses

Data were analyzed by unpaired t-test, 1-way ANOVA or mixed effect models for repeated measures (GraphPad Prism 8), to test the effect of TrkB agonist treatments with post-hoc multiple comparisons as appropriate. Statistical significance was set as p<0.05.

## 3 Results

### 3.1 LM22A-4 and TDP6 increase myelin sheath thickness during remyelination

We have previously shown treatment with TDP6, a structural mimetic of BDNF, enhances the number of axons remyelinated and increases myelin sheath thickness during recovery after 6-weeks cuprizone challenge in an oligodendroglial-TrkB dependent manner (Fletcher et al., 2018a). Here, we compared TDP6 with LM22A-4, a small molecule TrkB agonist reported to be a functional BDNF-mimetic (Massa et al., 2010). Demyelination by cuprizone feeding was confirmed by myelin basic protein (MBP) immunostaining, with severely reduced levels of MBP expression observed in animals taken at 6 weeks of cuprizone feeding (minimum = 2/cohort; Figure S2). Cuprizone feed was withdrawn and remaining animals received ICV minipumps containing aCSF (artificial cerebrospinal fliud), TDP6 (40µM) or LM22A-4 (500µM) for 7 days.

To examine the extent of remyelination, MBP-immunostaining in the caudal corpus callosum was assessed. This revealed both TDP6 and LM22A-4 treatment increased (p<0.0001) the percentage area of MBP^+^ staining compared to treatment with the aCSF vehicle (Fig. 1A, quantified in Fig. 1B). EM analysis indicated that mice treated with TDP6 exhibited a trend increase towards (p=0.09) more remyelinated axons compared to those receiving aCSF, whereas for those receiving LM22A-4, no increase was observed (p=0.46; Fig. 1C). Both TDP6 and LM22A-4 treatment resulted in a significant (p=0.002) reduction in mean g-ratio indicative of increased myelin thickness (Fig. 1D). Linear regression analysis of g-ratio against axon diameter (Fig. 1E) indicated that although both TDP6 and LM22A-4 treatments increase myelin sheath thickness during remyelination (Fig. 1D), TDP6 exerted a more consistent effect with a significant decrease in y-intercept (p=0.0032), but no change in slope (p=0.35) indicating that g-ratio was reduced across all axonal diameters, whereas for LM22A-4 there was a significant increase in slope (p=0.006), indicative of reduced g-ratio and thicker myelin on smaller diameter axons. Collectively, these data are consistent with our previous findings that BDNF-TrkB signaling increases myelin sheath thickness during remyelination *in vivo* (Fletcher et al., 2018a).

**Figure 1.**
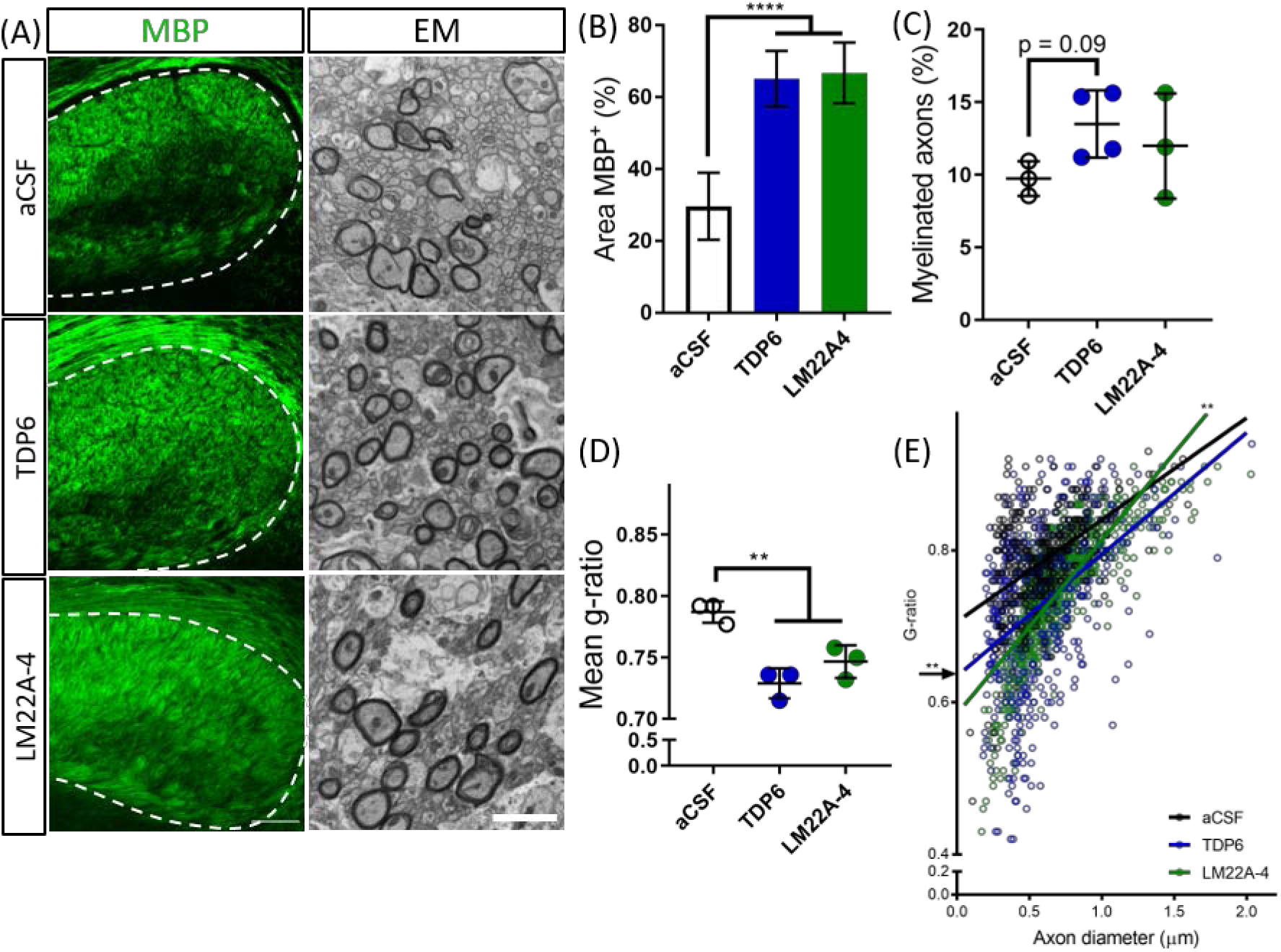
BDNF-structural mimetic, TDP6 and TrkB agonist, LM22A-4 increase myelin sheath thickness during remyelination in the cuprizone model of demyelination. (A) Representative MBP immunostaining (sagittal plane, scale bar=100µm) and electron micrographs (scale bar=2µm) of the caudal corpus callosum of aCSF vehicle, TDP6 and LM22A-4 treated cuprizone demyelinated mice. (B) Percentage of MBP staining increased (p<0.0001) in TDP6 and LM22A-4 treated mice compared to those that received aCSF vehicle (n=4-8/group). (C) The proportion of axons remyelinated trended towards increasing with TDP6 treatment (p=0.09) compared to aCSF, but was unchanged between aCSF and LM22A-4 (p=0.48) and LM22A-4 and TDP6 (p=0.31, n=3-4/group), however (D) the mean g-ratio was decreased (p=0.002) in both TDP6 and LM22A-4 treated mice, indicative of thicker myelin sheaths compared to those receiving aCSF vehicle (n=3/group). (E) Scatter plot of g-ratio against axonal diameter. Linear regression revealed that TDP6 treatment resulted in a decrease in y-intercept (p=0.0032) but no change in slope (p=0.35), while LM22A-4 treatment increased slope (p=0.0061), both indicative of increased myelin sheath thickness compared to aCSF treatment (n=3/group, min. 100 axons/animal). One-way ANOVA with Tukey’s post-hoc comparisons, p<0.05 considered significant. Mean ± SD plotted.

### 3.2 Treatment with LM22A-4 profoundly increases oligodendroglial densities during myelin repair

Next, we assessed oligodendroglial populations in the corpus callosum by co-immunostaining Olig2 with PDGFRα and CC1 to identify Olig2^+^PDGFRα^+^ oligodendrocyte progenitor cells (OPCs), Olig2^+^CC1^+^ post-mitotic oligodendrocytes and an Olig2^+^PDGFRα^−^CC1^−^ intermediate oligodendroglial population (Fig. 2A). Counts in the caudal corpus callosum revealed TDP6 and LM22A-4 increased the total population of Olig2^+^ oligodendroglia compared to treatment with aCSF vehicle (Fig. 2B, p<0.0001). Interestingly, LM22A-4 treatment exerted a more profound effect, increasing the density of Olig2^+^ oligodendroglia above TDP6 (Fig. 2B, p=0.0001). Both TDP6 and LM22A-4 treatments increased the density of Olig2^+^CC1^+^ post-mitotic oligodendrocytes compared to aCSF vehicle (Fig. 2D, p=0.013), consistent with the pro-differentiation effect of TrkB activation on oligodendroglia. However, assessment of Olig2^+^PDGFRα^+^ OPCs indicated LM22A-4 also increased the density of OPCs compared to treatment with TDP6 or aCSF vehicle (Fig. 2C, p=0.011). Overall, these data suggest that selectively targeting TrkB during remyelination primarily enhances oligodendroglial differentiation.

**Figure 2.**
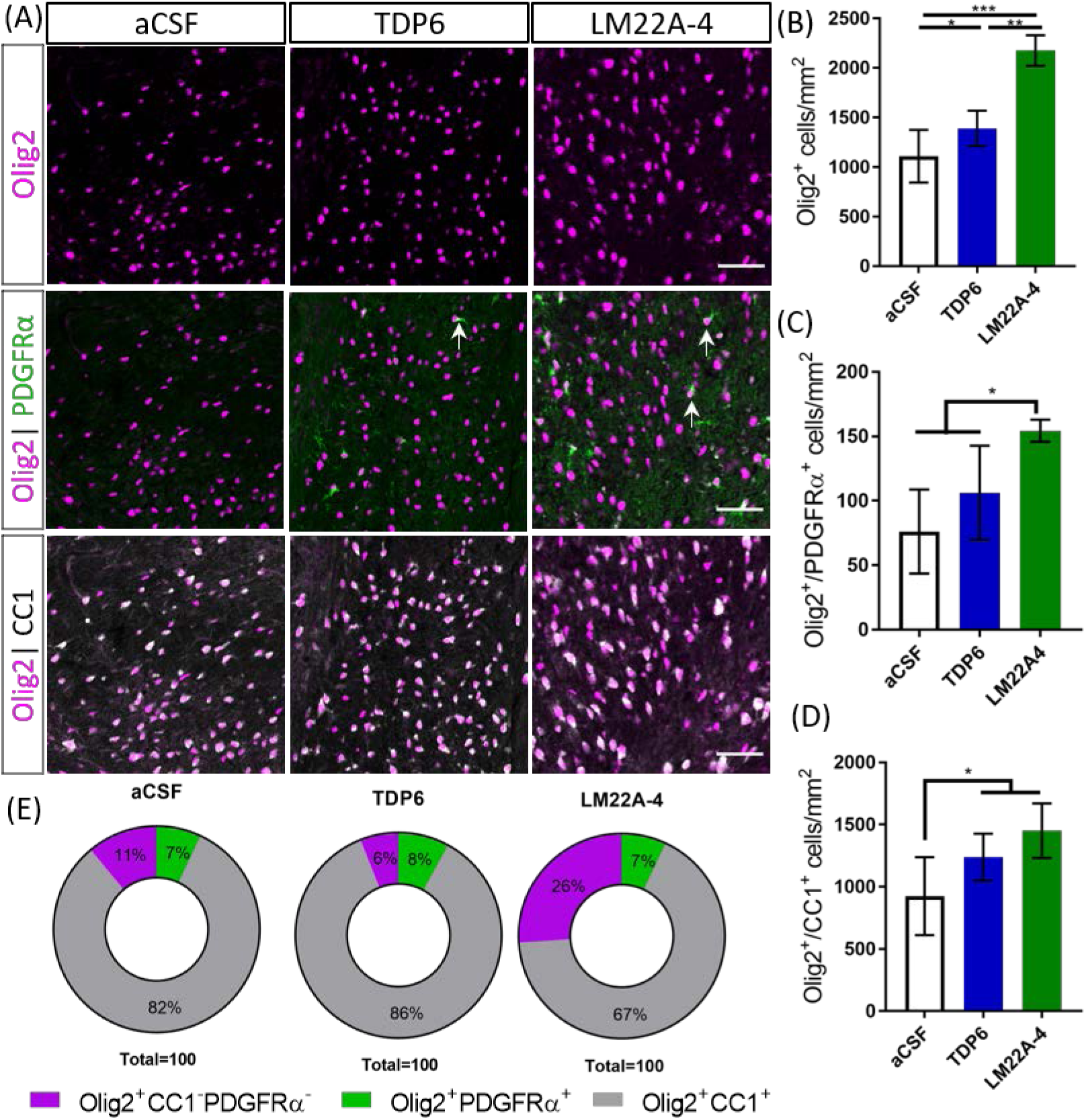
TDP6 and LM22A-4 increase oligodendroglial densities in the corpus callosum during remyelination in the cuprizone model of demyelination. (A) Representative micrographs of Olig2-CC1-PDGFRα immunostaining in the caudal corpus callosum of aCSF vehicle, TDP6 and LM22A-4 treated C57BL/6 mice. Arrows: Olig2^+^PDGFRα^+^ OPCs (sagittal plane, scale bar=50µm). (B) Olig2^+^ densities increased with LM22A-4 treatment (p<0.0001) compared to treatment with TDP6, which also increased Olig2^+^ cell density (p=0.048) compared to the aCSF vehicle (n = 4-8/group). (C) Olig2^+^PDGFRα^+^ OPC densities increased with LM22A-4 infusion (p=0.009) compared to the aCSF vehicle, but not compared to TDP6 treatment (p=0.12; n=4-8/group). (D) Both TDP6 (p=0.023) and LM22A-4 (p=0.007) infusions increased Olig2^+^CC1^+^ post-mitotic oligodendrocyte densities compared to the aCSF vehicle (n=4-8/group). (E) LM22A-4 treatment increased (p<0.0001) the proportion of Olig2^+^ only oligodendroglia compared to TDP6 and aCSF treatment (χ^2^ distribution test). For (A-D) one-way ANOVA with Tukey’s post-hoc comparisons, p<0.05 considered significant. Mean ± SD plotted.

To examine whether these effects of LM22A-4 and TDP6 were due to alterations in lineage progression during differentiation, the proportion of Olig2^+^PDGFRα^+^, Olig2^+^CC1^+^ and Olig2^+^ only cells out of the total Olig2^+^ population were assessed (Fig. 2E). The proportion of cells that were OPCs or post-mitotic oligodendrocytes were unchanged between groups. However, LM22A-4 treatment significantly increased the proportion of Olig2^+^ only cells (26±7%) compared to treatment with either TDP6 (6±8%) or aCSF vehicle (10±7%) (Fig 2E, mean ± SD, p<0.0001, χ^2^ distribution test). These data suggest LM22A-4 treatment may exert a greater effect than TDP6 to increase the proliferation or survival of oligodendroglia during myelin repair.

### 3.3 TrkB phosphorylation in the corpus callosum is elevated following treatment with TDP6 and LM22A-4 during remyelination

To determine if TDP6 and LM22A-4 infusions stimulated TrkB phosphorylation on oligodendroglia we performed triple immunolabelling for pTrkB^S478^ with PDGFRα and CC1 to identify OPCs and post-mitotic oligodendrocytes, respectively (Fig. 3A). This revealed that TDP6 and LM22A-4 infusions were successful, with an increased proportion of pTrkB^S478+^ cells in the corpus callosum during remyelination, compared to the aCSF vehicle (Fig. 3B, p=0.0022). Assessment of the proportion of pTrkB^S478+^ cells positive for the OPC marker PDGFRα indicated treatment with TDP6 and LM22A-4 had no effect on TrkB activation on OPCs (Fig. 3C, p=0.21). However, the proportion of TrkB^S478+^CC1^+^ post-mitotic oligodendrocytes increased with TDP6 treatment compared to treatment with LM22A-4 (Fig. 3D, p=0.046). These data suggest that LM22A-A can signal *via* TrkB during remyelination *in vivo* and is consistent with previous findings that TDP6 stimulates TrkB phosphorylation on CC1^+^ oligodendrocytes.

**Figure 3.**
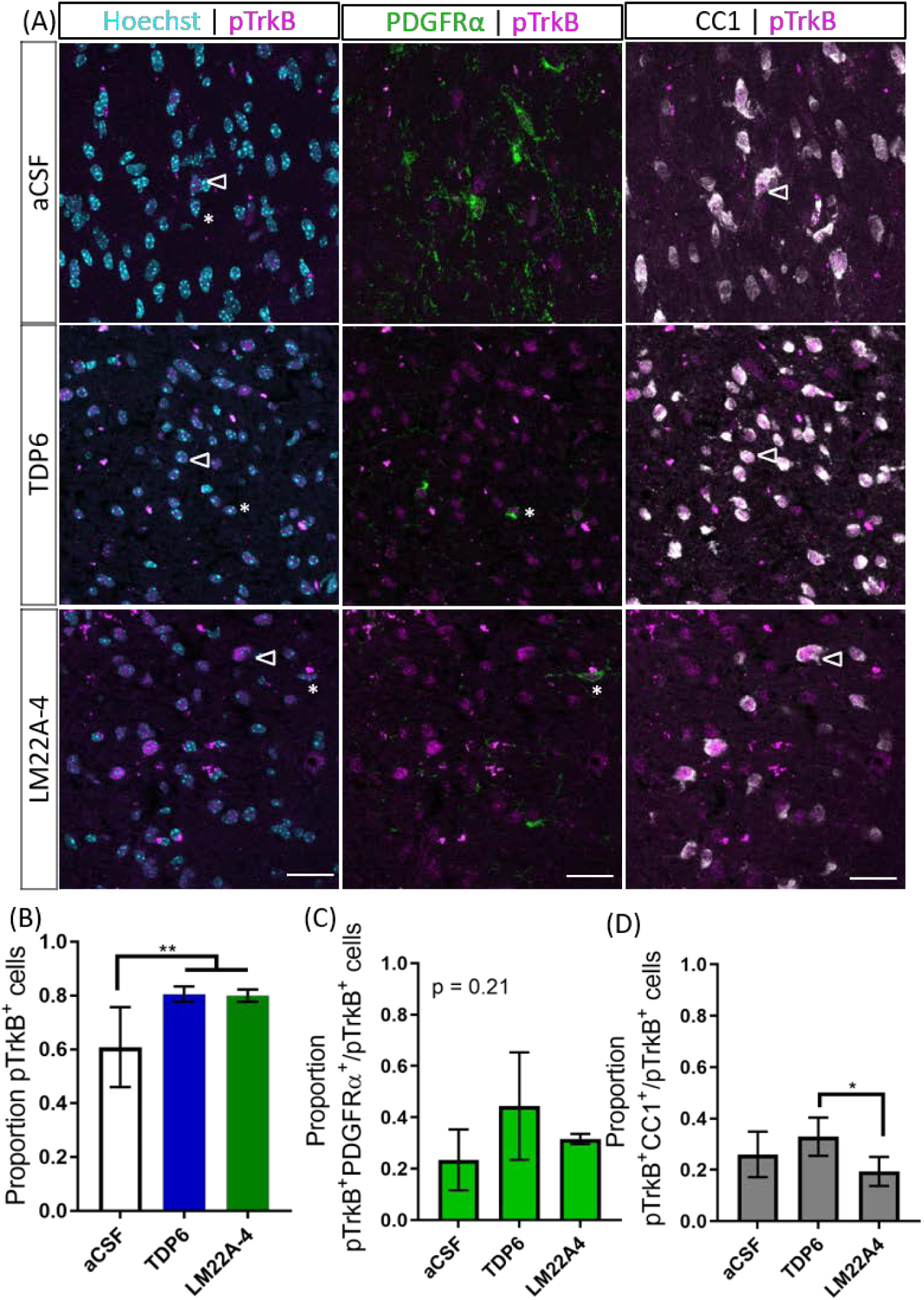
Phosphorylation of TrkB in the corpus callosum during remyelination was elevated following treatment with TDP6 and LM22A-4. (A) Representative micrographs of pTrkB^S478^-PDGFRα-CC1 immunostaining in the caudal corpus callosum of mice treated with aCSF vehicle, TDP6 or LM22A-4 (sagittal plane, scale bar=20µm). Asterisk: pTrkB^s478+^PDGFRα^+^ cell, open arrowhead: pTrkB^s478+^CC1^+^. (B) Proportion of pTrkB^s478+^ cells increased (p=0.0022) in the corpus callosum of mice treated with TDP6 and LM22A-4 compared to those that received aCSF vehicle (n=4-8/group). (C) The proportion of pTrkB^s478+^ cells also PDGFRα^+^ was unchanged across the three different treatments (p=0.21, n=4-8/group), while (D) the proportion of pTrkB^s478+^CC1^+^ cells increased (p=0.045) in mice that received TDP6 infusion compared to those receiving LM22A-4 (n=4-8/group). One-way ANOVA with Tukey’s post-hoc multiple comparisons, p<0.05 considered significant. Mean ± SD plotted.

### 3.4 LM22A-4 mediated increases in myelin sheath thickness and oligodendroglial densities require oligodendrocyte TrkB expression

To determine whether the effects of LM22A-4 on myelin sheath thickness and oligodendrocyte populations during myelin repair are dependent on oligodendroglial TrkB expression, we repeated the infusion experiment in CNPaseCre^+/−^ x TrkB^fl/fl^ mice in which TrkB is genetically deleted from maturing oligodendrocytes. These mice have a 3-fold reduction in TrkB^+^ oligodendroglia but adult myelination and oligodendrocyte populations are unaffected (Fletcher et al., 2018a). Cuprizone was administered for 6 weeks, and LM22A-4 or aCSF vehicle was infused via ICV minipumps for 7 days. Immunostaining for MBP revealed that LM22A-4 treatment in the oligodendroglial TrkB knockout mice had no effect on the percentage area of MBP^+^ immunostaining compared to the aCSF vehicle (Fig. 4A, quantified in Fig. 4B, p=0.21). Similarly, EM analysis (Fig. 4D) revealed there was no change in the proportion of axons myelinated with LM22A-4 treatment (Fig. 4C, p=0.85)) or the mean g-ratio (aCSF: 0.77 ± 0.087; LM22A-4: 0.78 ± 0.088, p=0.90, n=3-4/group, unpaired t-test). These data are consistent with oligodendroglial TrkB expression being necessary for LM22A-4 to increase myelin sheath thickness during remyelination.

**Figure 4.**
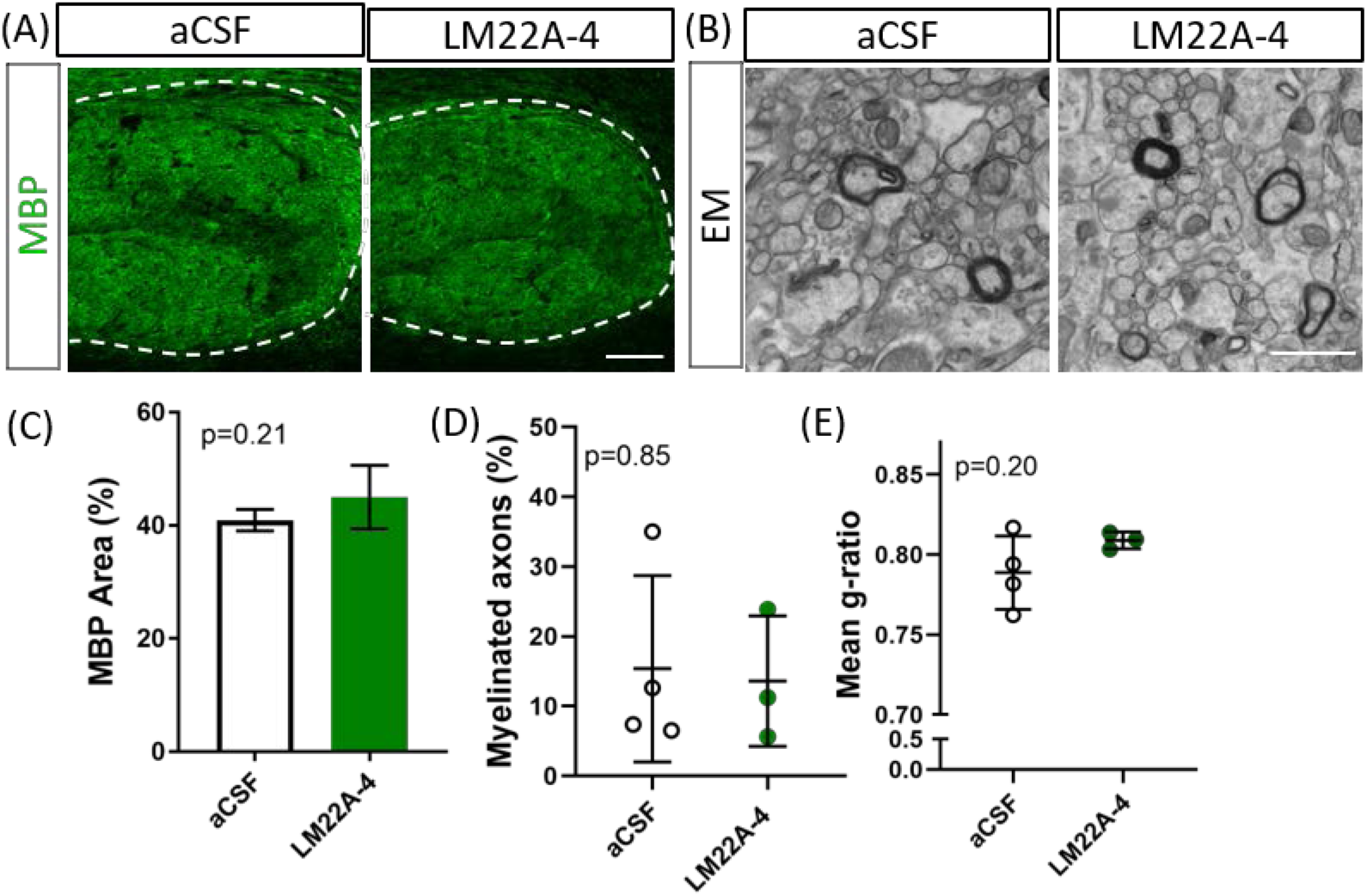
The effects of LM22A-4 on myelin sheath thickness during myelin repair require oligodendrocyte TrkB. (A) Representative micrographs of MBP immunostaining in the caudal corpus callosum of CNPaseCre^+/−^ x TrkB^fl/fl^ mice treated with aCSF vehicle or LM22A-4 (sagittal plane, scale bar=100µm). (B) Representative electron micrographs of the caudal corpus callosum of conditional TrkB knockout mice receiving aCSF vehicle or LM22A-4 (scale bar=2µm). (C) There was no change (p=0.21) in the percentage area of MBP^+^ immunostaining in the corpus callosi of oligodendroglial TrkB knockout mice treated with LM22A-4 compared to the aCSF vehicle. (D) The proportion of myelinated axons was unchanged between oligodendroglial TrkB knockout mice receiving aCSF vehicle and LM22A-4. (E) There was also no change in myelin sheath thickness as indicated by mean g-ratio, with LM22A-4 in CNPaseCre^+/−^ x TrkB^fl/fl^ mice. For (D-E) unpaired t-test with equal variance, p<0.05 considered significant, n=3-4/group. Mean ± SD plotted.

To determine if LM22A-4 treatment increased oligodendroglial populations during remyelination in oligodendroglial TrkB knockout mice, triple immunolabelling for Olig2-PDGFRα-CC1 was performed in the contralateral caudal corpus callosum (Fig. 5A). Counts revealed that LM22A-4 treatment had exerted no change in the density of Olig2^+^ oligodendroglia (p=0.91; Fig. 5B), Olig2^+^PDGFRα^+^ OPCs (p=0.38; Fig 5C) or Olig2^+^CC1^+^ post-mitotic oligodendrocytes (p=0.94; Fig. 5D) compared to the aCSF vehicle. Similarly, there was no difference in the proportion of Olig2^+^ only cells between aCSF vehicle and LM22A-4 treatment in the oligodendroglial TrkB knockout mice (Fig. 5E, p=0.33, χ^2^ distribution test). Collectively, these data confirm that the action of LM22A-4 in increasing oligodendroglial populations is dependent on oligodendroglial expressed TrkB.

**Figure 5.**
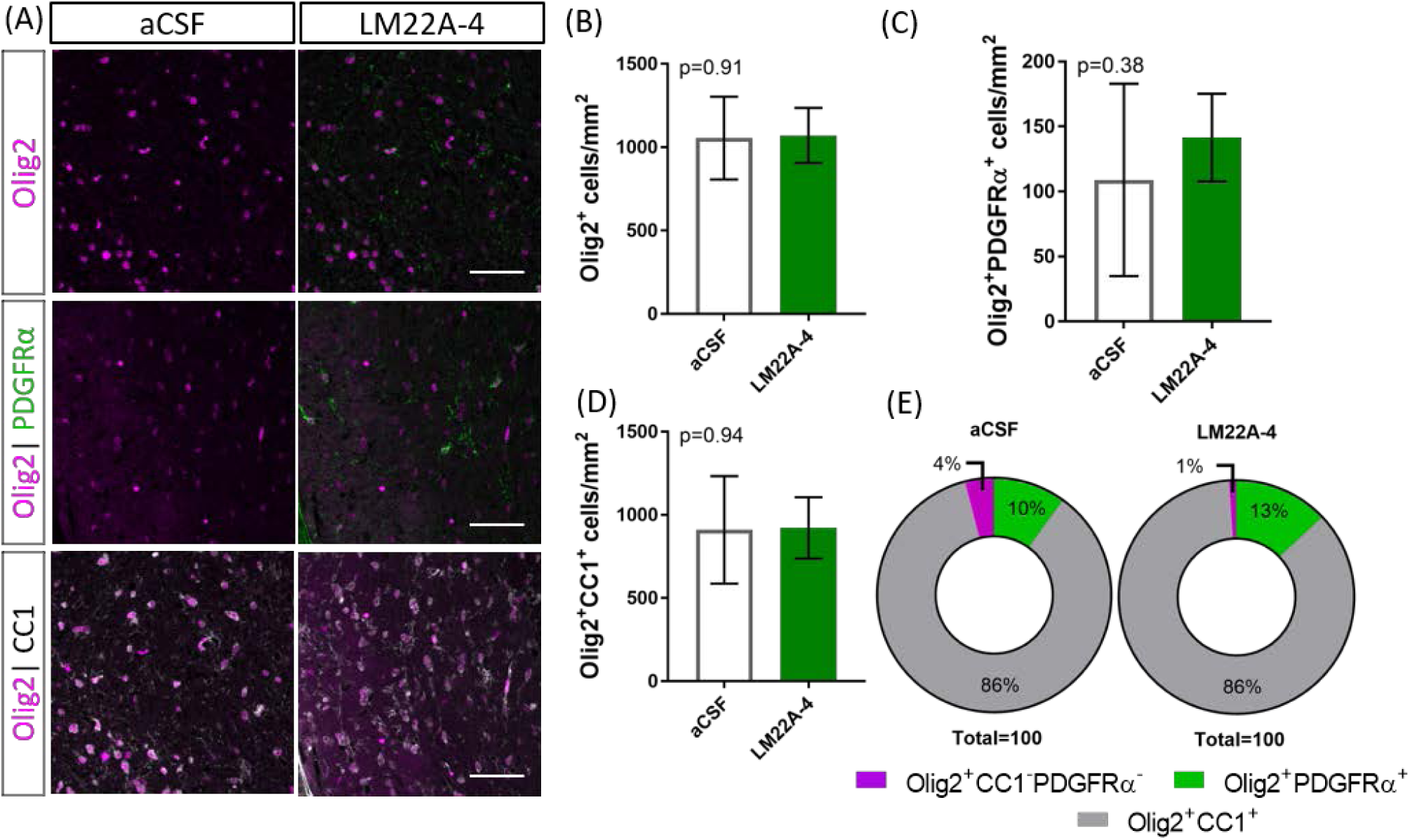
Increased oligodendroglial density mediated by LM22A-4 during myelin repair requires oligodendroglial TrkB. (A) Representative micrographs of Olig2-CC1-PDGFRα immunostaining in the caudal corpus callosum of CNPaseCre^+/−^ x TrkB^fl/fl^ mice treated aCSF vehicle or LM22A44 (sagittal plane, scale bar=20µm). (B) Density of Olig2^+^ oligodendroglia was unchanged (p=0.91) in TrkB conditional knockout mice treated with LM22A-4 compared to aCSF vehicle. (C) Olig2^+^PDGFRα^+^ OPC densities were unchanged (p=0.38) in oligodendroglial TrkB knockout mice treated with LM22A-4 compared to aCSF vehicle. (D) Density of Olig2^+^CC1^+^ oligodendrocytes in TrkB conditional knockout mice was unchanged (p=0.94) with LM22A-4 infusion compared to aCSF vehicle. (E) There was no change (p=0.34) in the proportion of oligodendroglia that were Olig2^+^ only, Olig2^+^PDGFRα^+^ or Olig2^+^CC1^+^ with LM22A-4 or a CSF vehicle treatment (χ^2^ distribution test). For (A-D) unpaired t-test with equal variance, p<0.05 considered significant, n=3-4/group. Mean ± SD plotted.

To examine TrkB phosphorylation in LM22A-4 treated oligodendroglial TrkB knockout mice, immunohistochemistry for pTrkB^S478^ with oligodendrocyte markers PDGFRα and CC1 was performed (Fig. 6A). Analysis of the caudal corpus callosum revealed that in the oligodendroglial TrkB knockout mice LM22A-4 treatment did not increase the proportion of pTrkB^S478+^ cells compared to the aCSF vehicle (Fig. 6B, p=0.24). This was also reflected with no change in the proportion of pTrkB^S478+^ cells positive for oligodendroglial markers PDGFRα (p=0.99; Fig. 6C) or CC1 (p>0.99; Fig. 6D). These data indicate that for LM22A-4 mediated TrkB phosphorylation during remyelination requires oligodendroglial TrkB expression.

**Figure 6.**
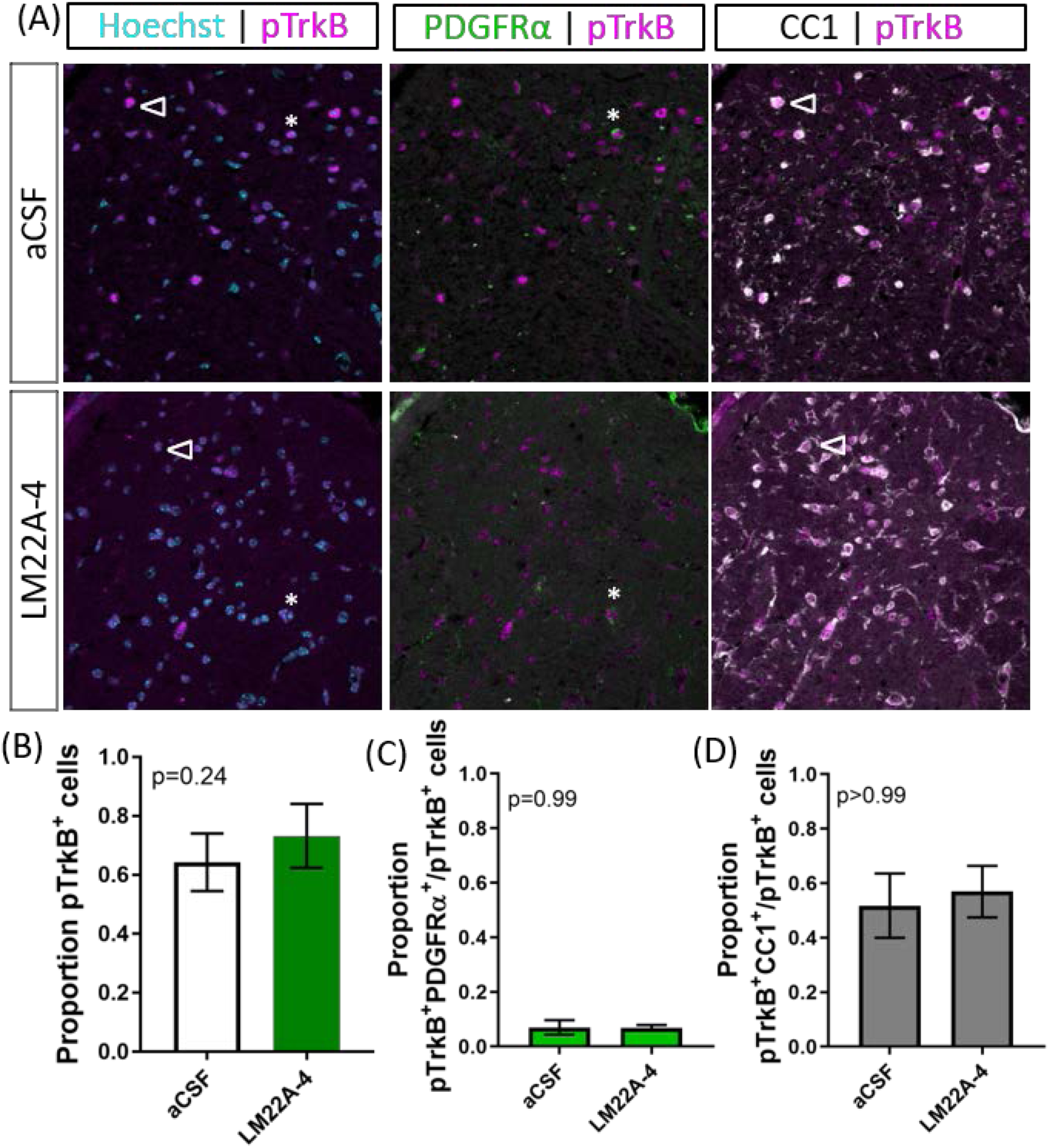
TrkB phosphorylation with LM22A-4 treatment does not increase in the corpus callosi of mice with oligodendroglial TrkB deleted. (A) Representative micrographs of pTrkB^S478^-PDGFRα-CC1 immunostaining in the corpus callosum of CNPaseCre^+/−^ x TrkB^fl/fl^ mice treated with LM22A-4 or aCSF vehicle (sagittal plane, scale bar=20µm). (B) Proportion of pTrkB^s478+^ cells was unchanged (p=0.24) in oligodendroglial TrkB knockout mice treated with LM22A-4 compared to aCSF vehicle. Similarly, (B) the proportions of pTrkB^s478+^ PDGFRα^+^ and (C) pTrkB^s478+^CC1^+^ cells were not changed (p=0.99, p>0.99 respectively) in conditional knockout mice treated LM22A-4 compared to those that received aCSF vehicle. Unpaired t-test with equal variance, p<0.05 considered significant, n=3-4/group. Mean ± SD plotted.

### 3.5 TrkB signaling dynamics initiated by LM22A-4 do not mimic BDNF

To determine if LM22A-4 elicits a signaling cascade mimicking typical BDNF-TrkB signaling, we generated an isogenic stable TrkB expressing HEK293 (293-TrkB) cell line using the Flp-In system (Supplementary Fig. 1A). TrkB expression in the 293-TrkB cells was confirmed by Western blot and compared to TrkB expression generated by transiently transfecting Flp-In HEK293 cells with the Ntrk2 expression vector. This revealed that transiently transfected cells overexpress both mature glycosylated and unprocessed TrkB receptors, whereas the 293-TrkB cells express only the fully mature glycosylated form (Supplementary Fig. 1B). To confirm that the 293-TrkB cells responded to BDNF, cells were treated with BDNF (0.04nM to 40nM) for 15 mins (Supplementary Fig. 1C) which resulted in increasing levels of TrkB and ERK1/2 phosphorylation (Supplementary Fig. 1D).

As determined in the original report characterizing LM22A-4 as functional BDNF mimetic (Massa et al., 2010), we used 500nM as the standard concentration for our *in vitro* studies. The 293-TrkB cells were treated with 4nM BDNF or 500nM LM22A-4 for a time course of 5, 15, 30, 60 and 240mins and assessed for TrkB and ERK1/2 phosphorylation by Western blot (Fig. 7A). Densitometric analysis (Fig. 7B) revealed that compared to BDNF treatment, which increased TrkB phosphorylation within 5mins (p=0.012), LM22A-4 did not significantly increase levels of phosphorylated TrkB until 240mins of treatment (p=0.02). The effects of LM22A-4 treatment on ERK1/2 phosphorylation where levels peaked at 5mins of treatment, and significantly declined compared to BDNF from 15 to 240mins (Fig. 7C). Collectively, these data indicate that LM22A-4 does not elicit a signaling cascade that mimics typical BDNF-TrkB signaling, in particular suggesting that pERK1/2 is upstream of TrkB phosphorylation in the pathway stimulated by LM22A-4.

**Figure 7.**
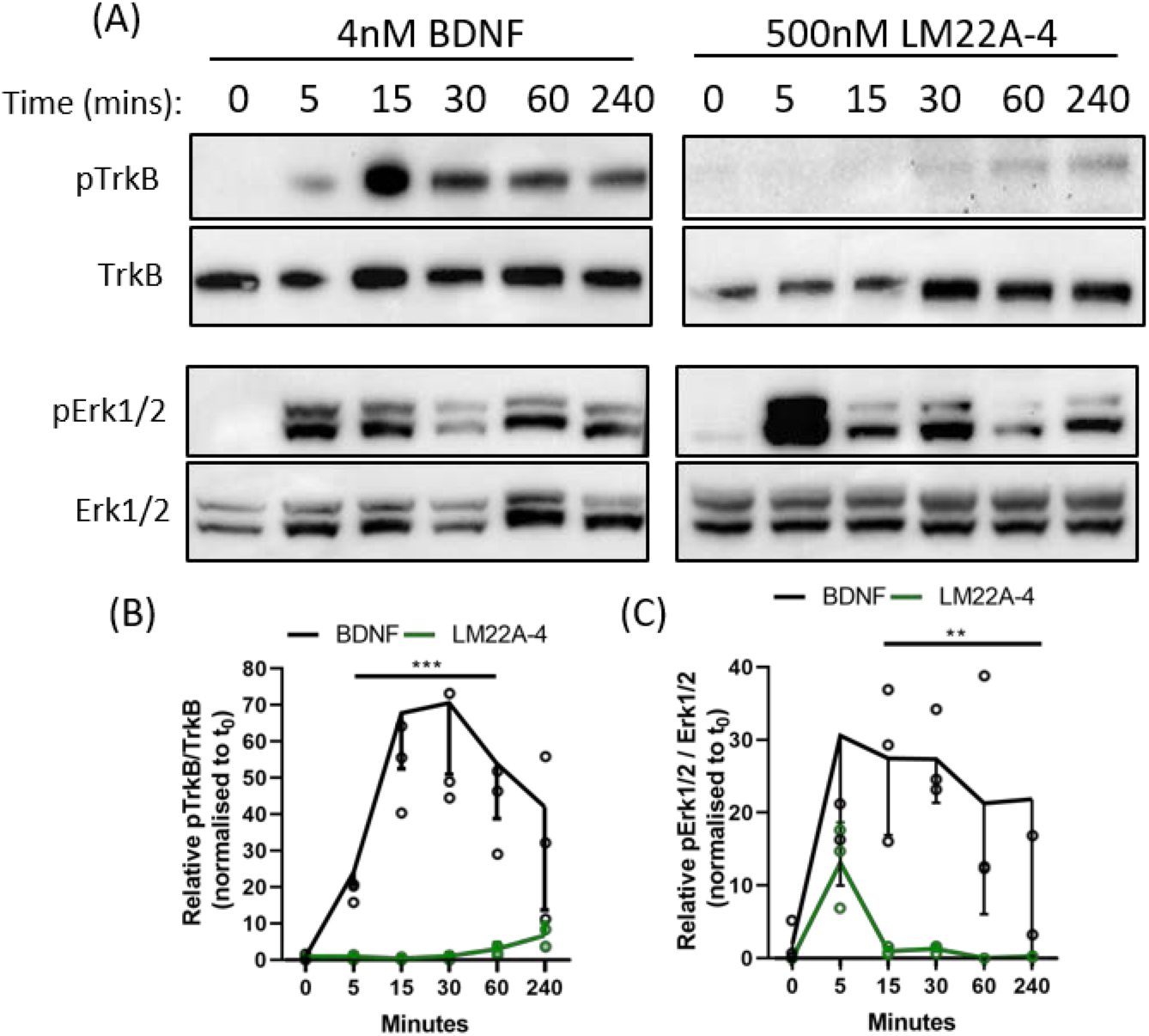
Dynamics of TrkB signaling initiated by LM22A-4 do not mimic BDNF in a stably expressing TrkB isogenic HEK293 cell line. (A) Representative western blots of isogenic TrkB FlpIn HEK293 cell lysates treated with 4nM BDNF or 500nM LM22A-4 over a time course of 0, 5, 15, 30, 60 and 240 mins. Densiometric analysis of western blots revealed that (B) BDNF elevated (p=0.0003) levels of phosphorylated TrkB from 5mins until 240mins when it returned to similar levels elicited by LM22A-4 treatment, which did not increase over time (p=0.10). (C) Levels of Erk1/2 phosphorylation increased (p=0.002) after 5mins of BDNF treatment was sustained until 240mins, while phosphorylated Erk1/2 levels mediated by LM22A-4 treatment were similar to those evoked by BDNF at 5mins (p=0.051), but elevated levels were not sustained. Mixed effects model, fixed effects: treatment and time, random effects: plate, *n*=3 independent cultures, p<0.05 considered significant. Mean ± SD plotted.

## 4 Discussion

There is an unmet clinical need for therapies that promote myelin repair to halt disease progression in MS. Here, we have shown that targeting oligodendroglial TrkB activation during remyelination increases post-mitotic oligodendrocyte density and myelin sheath thickness. By comparing TDP6, a structural peptide mimic of the Loop 2 region of BDNF, and LM22A-4, a small molecule TrkB agonist we have identified that while both promote myelin repair, they exert disparate effects upon OPCs and intermediate oligodendroglial populations. The effects of both TDP6 and LM22A-4 are dependent on the expression of oligodendroglial TrkB, indicating that while these two molecules putatively activate the same receptor, they may result in biased or differential signaling within oligodendroglia, as indicated by the differences in remyelination profile and progression of oligodendrocyte differentiation. This may reflect our *in vitro* analysis comparing the effect of LM22A-4 and BDNF upon TrkB and ERK1/2 phosphorylation which demonstrates that LM22A-4 activates BDNF-TrkB signaling pathways with substantially different kinetics and magnitude, compared to BDNF. Importantly, use of the oligodendroglial TrkB knockout mice to test its therapeutic efficacy provides the first *in vivo* genetic evidence that LM22A-4 action requires the presence of TrkB, although our data suggests that the mode of TrkB activation may differ to what was previously appreciated.

For decades the therapeutic promise of BDNF-TrkB signaling as a treatment for neurodegenerative and demyelination conditions has been recognized (Fletcher et al., 2018a; Longo and Massa, 2013; McTigue et al., 1998). However, the poor pharmacokinetic properties of BDNF have led to focused development of BDNF mimetics and small molecule TrkB agonists, including TDP6 and LM22A-4 among others (Boltaev et al., 2017; Longo and Massa, 2013). Previously, we showed that the structural BDNF-mimetic TDP6 enhances remyelination, increasing the proportion of axons remyelinated, and density of post-mitotic oligodendrocytes compared to treatment with the vehicle and BDNF (Fletcher et al., 2018a). We found that LM22A-4 increased oligodendroglial density to a greater degree than TDP6, but instead of solely affecting maturing cells, LM22A-4 also increased OPC density. Both these effects were dependent on oligodendroglial TrkB expression. This raises the ongoing, unresolved question of whether oligodendroglial TrkB signaling exerts a direct influence on oligodendroglial proliferation and survival during remyelination, in addition to its well-established pro-differentiation effect (Fletcher et al., 2018a; Goebbels et al., 2017; Xiao et al., 2011). Previous studies in the BDNF heterozygous global knockout mice showed that oligodendroglial populations are sensitive to low BDNF levels during cuprizone demyelination, with reduced proliferating OPCs and subsequently differentiated oligodendrocytes (Tsiperson et al., 2015; VonDran et al., 2011). This appears to contrast our observations, where exogenous BDNF or TDP6 exerted no effect on the density or proliferative fraction of OPCs during remyelination after cuprizone (Fletcher et al., 2018a). The different observations between these two distinct experimental approaches may ultimately reflect context, wherein oligoendroglia subjected to a lifetime of BDNF haploinsufficiency simply behave differently.

The selective influence that LM22A-4 exerted upon OPCs remains to be explained. The doses used for TDP6 and LM22A-4 were determined from reported concentrations required for these compounds to mimic the neurotrophic activity of BDNF in *in vitro* myelinating co-culture or neuronal survival assays (Massa et al., 2010; O’Leary and Hughes, 2003; Wong et al., 2014). TDP6, like BDNF, demonstrates a strong signaling bias for MAPK/ERK in oligodendrocytes (Du et al., 2006; Wong et al., 2014; Xiao et al., 2011). In contrast, a recent report in a rat traumatic epileptogenesis model indicates LM22A-4 may demonstrate bias towards PI3K/Akt signaling (Gu et al., 2018). The PI3K/Akt and MAPK/ERK signaling pathways are known to act independently and cooperatively in oligodendrocytes to regulate distinct stages of oligodendrocyte myelination (Dai et al., 2014; Ishii et al., 2019) and PI3K/Akt signaling has been identified as necessary for OPC survival *in vitro* (Ebner et al., 2000; Ness et al., 2002). The potential signal bias towards PI3K/Akt over MAPK/ERK may explain why LM22A-4 elicited an increase in OPCs, as well as the anticipated increase in post-mitotic oligodendrocyte densities.

Both TDP6 and LM22A-4 infusion increased myelin sheath thickness in a manner dependent on oligodendroglial TrkB expression, consistent with the demonstrated effects of TrkB signaling *via* MAPK/ERK in oligodendrocytes to promote myelin sheath growth (Ishii et al., 2012, 2016). It also confirms our previous findings where infusion of exogenous BDNF or TDP6 increased myelin sheath thickness during remyelination (Fletcher et al., 2018a). Intriguingly, LM22A-4 exerted its effect on myelin thickness almost selectively on smaller diameter axons, which was completely abrogated in the oligodendroglial TrkB knockout mice. This contrasts with TDP6, and our previous findings with BDNF (Fletcher et al., 2018a), where myelin sheath thickness increased across all axonal diameters. However, it echoes our findings that TDP6 treatment in oligodendroglial TrkB knockout mice resulted in increased myelination of small diameter axons during myelin repair (Fletcher et al., 2018a). It is tempting to speculate that small diameter axons are exerting a selective effect in both instances, but it is critical to distinguish growth in myelin thickness as an oligodendrocyte-driven function (Ishii et al., 2012). To date, a direct axonal signal that instructs oligodendrocytes to increase myelin thickness has not been identified, although the number of myelin wraps is known to increase as circuit activity increases with the maturing brain (Sturrock, 1980). In contrast, initiation of myelination, particularly for small diameter axons is known to require axonally derived signals (Bechler et al., 2017; Gautier et al., 2015), suggestive that TrkB expression by neurons may potentially confer a pro-myelinating signal to oligodendrocytes.

Concerningly, a recent report indicates that LM22A-4 does not activate TrkB at all (Boltaev et al., 2017). The fact that LM22A-4 failed to promote remyelination and increase oligodendroglial density in the oligodendrocyte TrkB knockout mice clearly indicates TrkB is necessary for the action of LM22A-4 and this is the first *in vivo* genetic evidence that LM22A-4 requires TrkB for activity. Although LM22A-4 may exert its effect through direct activity on TrkB, our data and findings by Boltaev et al. (2017) raise the possibility that LM22A-4 is exerting an indirect effect, potentially by increasing BDNF or NT-4 expression in the demyelinated lesion. This is a possibility we certainly cannot discount. It is important to note, however, that Boltaev et al. (2017) used a model system of cortical cultures to assess TrkB activation. Our observations in the 293-TrkB cells identified that LM22A-4 produced a spike of ERK1/2 phosphorylation after 5mins independent of any evidence of TrkB phosphorylation, but then identified TrkB phosphorylation after 4 hours of LM22A-4 exposure. This supports a model in which TrkB activation is an event downstream of LM22A-4 activity at another receptor, although this also contrasts with findings from the original report wherein TrkB phosphorylation was detected in cultured hippocampal neurons within 60mins (Massa et al., 2010). Notably, ERK1/2 phosphorylation was detected in these cultures within 10mins (Massa et al., 2010). Whilst 293-TrkB cells and the two types of neuronal cultures are contextually quite different cells, Boltaev et al. (2017) limited LM22A-4 treatment to 2 hours, potentially missing this delayed response. Such a delay in Trk phosphorylation has been reported previously (Lee and Chao, 2001) and is a pattern consistent with Trk-receptor transactivation.

The extended 4-hour timeframe required to detect TrkB phosphorylation following LM22A-4 treatment *in vitro* is consistent with Trk-receptor transactivation, where it can take up to 6 hours to elicit detectable Trk receptor phosphorylation, and results in signal bias towards Akt (Lee and Chao, 2001). We propose that LM22A-4 is mediates its increase in OPC density during remyelination by Trk-transactivation potentially *via* GPCRs (Fig. 8). Multiple GPCRs are known to be critical for OPC proliferation and regulating OPC differentiation towards myelination (Chen et al., 2009; Giera et al., 2015; Yang et al., 2016). In our model, we hypothesise that LM22A-4 acts *via* unidentified GPCRs which engage Src-family kinases, most likely Fyn in oligodendrocytes; this results in intracellular phosphorylation of TrkB receptors confined in transport vesicles and not expressed at the cell surface (Fig. 8). This could be tested *in vitro* with LM22A-4 treated TrkB-293 cells or primary oligodendrocytes co-treated with Src inhibitors. Full understanding of the mode of action for LM22A-4 is warranted in order to optimize its therapeutic potential. Overall, our hypothesised mode of action for LM22A-4 is parsimonious with known roles of Src-family kinases / Fyn in Trk-transactivation (Rajagopal et al., 2004; Rajagopal and Chao, 2006) and oligodendroglial function (Colognato et al., 2004; Peckham et al., 2016; Sperber et al., 2001), as well as our current findings of delayed TrkB phosphorylation *in vitro* and TrkB-dependent remyelination outcomes *in vivo*.

**Figure 8.**
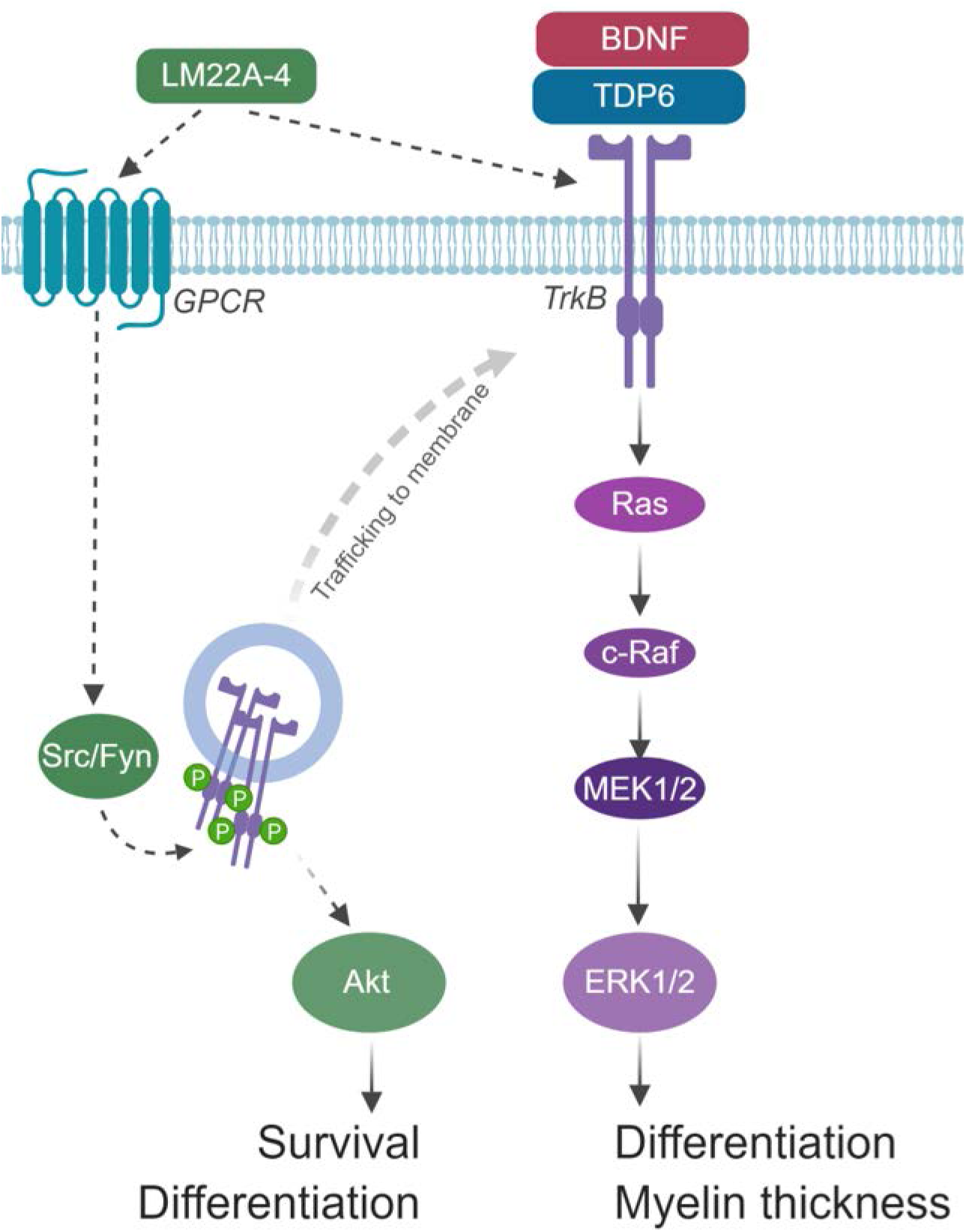
Schematic of hypothesized mode of action for LM22A-4 in promoting remyelination. LM22A-4 acts as a ligand to directly activate an unidentified GPCR, which initiates Src-family kinase, most likely Fyn, activation. Src-family members regulate the phosphorylation of Trk receptors on intracellular membranes, initiating Trk-specific signaling potentially biased towards PI3K/Akt, which regulates oligodendrocyte survival and differentiation. Activated Trk receptors may subsequently be trafficked to the cell membrane to initiate typical Trk signaling. Figure made in ©BioRender – biorender.com)

We have demonstrated targeting TrkB activation on oligodendrocytes either *via* a peptide mimetic or small molecule partial agonist, enhances myelin repair after a central demyelinating insult by increasing the density of post-mitotic oligodendrocytes and increasing myelin sheath thickness. By directly comparing these two strategies we have also provided necessary insight on the recent controversies about the fidelity of small molecule TrkB agonists. Through *in vivo* genetic deletion of oligodendroglial TrkB we have shown LM22A-4 is dependent on TrkB receptor expression for its effects on increasing oligodendroglial populations and myelin thickness, while our *in vitro* studies have shown that LM22A-4 acts in delayed manner, potentially through GPCR-mediated TrkB transactivation. Overall, our results further verify that targeting TrkB with small molecule mimetics is a viable therapeutic strategy to promote myelin repair in central demyelinating diseases, such as MS.

## 6 Contribution to the Field Statement

Multiple sclerosis (MS) is caused by autoimmune attack against the myelin sheath that results in sensorimotor, cognitive and psychosocial dysfunction. Critically, the brain’s innate capacity for myelin repair is also impaired in MS, contributing to chronic axonal damage and nerve cell death. To stop disease progression, there is an urgent unmet clinical need for remyelinating therapies. Here, we evaluate two strategies to mimic the pro-myelinating effect of brain-derived neurotrophic factor (BDNF) by targeting its TrkB receptor on myelin producing cells in a preclinical model of MS. We show that both a structural peptide-mimetic and a small molecule TrkB agonist enhance myelin repair, after a demyelinating insult by increasing the density of myelinating cells and increasing myelin sheath thickness. Importantly, we also resolve outstanding issues in the field regarding the fidelity of partial TrkB agonist, LM22A-4. We show that the *in vivo* effects on myelin repair mediated by LM22A-4 are dependent on oligodendrocyte expressed TrkB. However, the timeframe required for TrkB phosphorylation to occur is extended and reminiscent of Trk-receptor transactivation. Overall, our findings shed light on LM22A-4 mechanism of action and provide support for targeting TrkB activation as a therapeutic to promote myelin repair in central demyelinating disease.

## 7 Author Contributions

JF, JX and SM conceived and designed the study. JF, HN, RW, AP and SM performed experiments. JF, HN and SF analysed data. SF provided reagents/analytic tools. JF wrote the first draft. JF, HN, SF, JX and SM reviewed and revised the manuscript. All authors read, revised and approved the submitted version.

## 9 Acknowledgments

The authors acknowledge the staff and facilities of the Biological Optical Microscope Platform, The University of Melbourne and the Centre for Advanced Microscopy and Histology, Peter MacCallum Centre.

## 10 Supplementary Material

Supplementary Figure 1. Generation of isogenic TrkB expressing Flp-In HEK293 (TrkB-293) cells

Supplementary Figure 2: MBP immunostaining to confirm successful demyelination with 6 weeks’ cuprizone feeding

## 11 Data Availability Statement

The raw data supporting the conclusions of this manuscript will be made available by the authors, without undue reservation, to any qualified researcher.

