## Supplemental Figures 1 and 2 for "TrkB Agonist LM22A-4 Increases Oligodendroglial Populations During Myelin Repair in the Brain"

### *Supplementary Material*

#### 1 Supplementary Figures

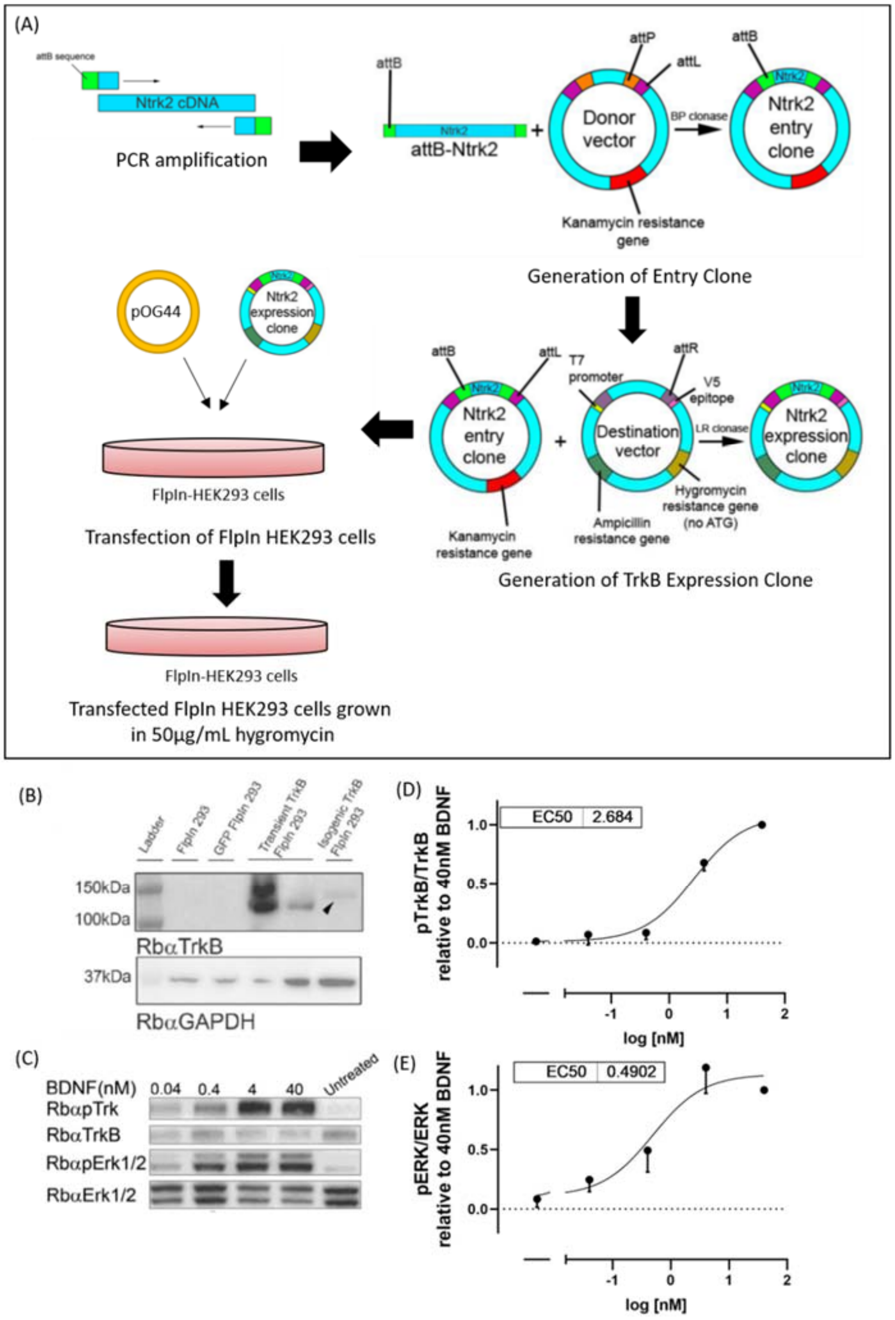


**Supplementary Figure 1.** (A) Schematic of the generation of isogenic TrkB-293 cells using the Flp-In system. (B) Representative Western blot demonstrating that TrkB-293 cells stably express the mature glycosylated form of TrkB. (C) Representative Western blot showing that (D) TrkB and (E) ERK1/2 phosphorylation in TrkB-293 cells demonstrate a dose response to BDNF with an $EC_{50}$ of 2.7nM and 0.5nM BDNF respectively.

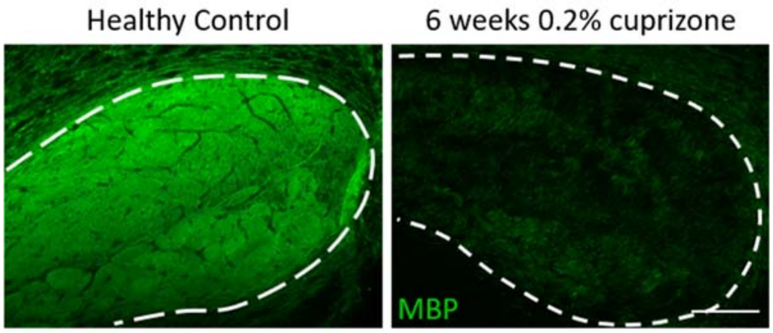


**Supplementary Figure 2.** Successful demyelination in the corpus callosum was confirmed with immunostaining for myelin basic protein (MBP). Min. $n$=2/cohort.
